# No evidence for basigin/CD147 as a direct SARS-CoV-2 spike binding receptor

**DOI:** 10.1101/2020.07.25.221036

**Authors:** Jarrod Shilts, Gavin J. Wright

## Abstract

The spike protein of SARS-CoV-2 is known to enable viral invasion into human cells through direct binding to host receptors including ACE2. An alternate entry receptor for the virus was recently proposed to be basigin/CD147. These early studies have already prompted a clinical trial and multiple published hypotheses of the role of this host receptor in viral infection and pathogenesis. We sought to independently characterize the basigin-spike protein interaction. After conducting several lines of experiments, we report that we are unable to find evidence supporting the role of basigin as a putative spike-binding receptor. Recombinant forms of both the entire ectodomain and S1 domain of the SARS-CoV-2 spike protein that directly bind ACE2 do not interact with basigin expressed on the surface of human cells. Using specialized assays tailored to detect receptor interactions as weak or weaker than the proposed basigin-spike binding, we report no evidence for direct binding of the viral spike to either of the two common isoforms of basigin. Given the pressing need for clarity on which targets of SARS-CoV-2 may lead to promising therapeutics, we present these findings to allow more informed decisions about the translational relevance of this putative mechanism in the race to understand and treat COVID-19.

## Introduction

The sudden emergence of SARS-CoV-2 in late 2019 has demanded extensive research be directed to resolve the many unknown aspects of this previously-unknown virus. One essential question is what host factors the virus uses to recognize and invade human cells. SARS-CoV-2, as with other members of the coronavirus family, invades host cells using the large trimeric spike proteins on their surfaces. A series of studies published within the first months of the COVID-19 pandemic independently confirmed that the same angiotensin-converting enzyme 2 (ACE2) receptor that was found to mediate SARS spike binding to human cells also mediates SARS-CoV-2 binding to human cells^1–3^. However, for previous coronaviruses closely related to SARS-CoV-2 including SARS and MERS, multiple different host receptors have been described with roles facilitating viral invasion^4–7^, making it plausible that additional interaction partners for the SARS-CoV-2 spike may remain undiscovered. Among the most prominent claims for an alternate SARS-CoV-2 host receptor comes from a report identifying basigin (CD147) as a binding partner for the SARS-CoV-2 spike protein with functional significance in viral invasion^9^. Based on a previously-published indirect interaction between the SARS spike protein and cyclophilin A for which basigin appeared to be involved, basigin was subsequently found to directly bind the spike SARS-CoV-2 spike protein with reasonably high affinity (185 nM, compared to 5-20 nM reported for the similarly high-affinity spike-ACE2 binding^2,10^). Direct binding between the SARS-CoV-2 spike “receptor-binding region” of the S1 domain and basigin was demonstrated in those reports by co-immunoprecipitation, surface plasmon resonance, and enzyme-linked immunosorbent assays (ELISAs).

Notably, the original finding that basigin is a possible alternative SARS-CoV-2 receptor has already translated into an open-label clinical trial of a humanized therapeutic monoclonal antibody against basigin, meplazumab, which reported striking improvements in COVID-19 patients treated with antibody^8^. Basigin represents an attractive medical target because therapeutic agents have already been developed that target basigin based on basigin’s previously-established role as an essential host receptor for invasion of the malaria parasite *Plasmodium falciparum*^11,12^. The claim that basigin acts as a host receptor for SARS-CoV-2 has already featured in published articles discussing the prioritization of therapeutics^13^, and has been the subject of published analyses looking at basigin expression on the assumption that it serves as a viral entry factor^14–16^.

We sought to validate the interaction between the SARS-CoV-2 spike protein and human basigin after observing that the result had not yet been reproduced despite intense interest in the interaction’s proposed consequences. Using a variety of sensitive approaches for detecting binding interactions and validated reagents, we were unable to find any supporting evidence for a direct interaction of the SARS-CoV-2 spike protein with basigin. Based on our findings, we encourage caution in approaches aimed at addressing the current pandemic caused by SARS-CoV-2 which are rooted in the assumption that basigin acts as a viral recognition receptor without further evidence.

## Results

We first investigated whether basigin (BSG) expressed on the surface of human cell lines could bind the spike protein of SARS-CoV-2. The previous reports of this interaction had not performed any binding experiments on full-length basigin displayed on the surface of cells^9^. First, we synthesized constructs to recombinantly express the spike protein of SARS-CoV-2. We emulated the published designs of spike constructs previously determined to be folded and functional^17^. Using a mammalian HEK293 expression system to increase the chances that preserve structurally-critical post-translational modifications would be preserved^18^, we produced both the full extracellular domain of the spike protein, and the S1 domain of the spike that mediates all known receptor binding events (Figure 1A). When HEK293 cells were transiently transfected with cDNA overexpression plasmids for ACE2, the transfected cells became strongly stained by fluorescent tetramers of spike protein in either S1 and full forms (Figure 1B). However, no similar gain of binding was observed with HEK293 cells transfected with BSG cDNA. We also noted that these HEK293 cell lines express BSG at high levels even without cDNA overexpression (Figure 1C), yet despite this spike protein tetramers had no detectable background staining of our HEK293 cells without ACE2 in either our experiment or similar experiments reported with SARS-CoV-2 and HEK293 cells^19–21^.

**Figure 1.**
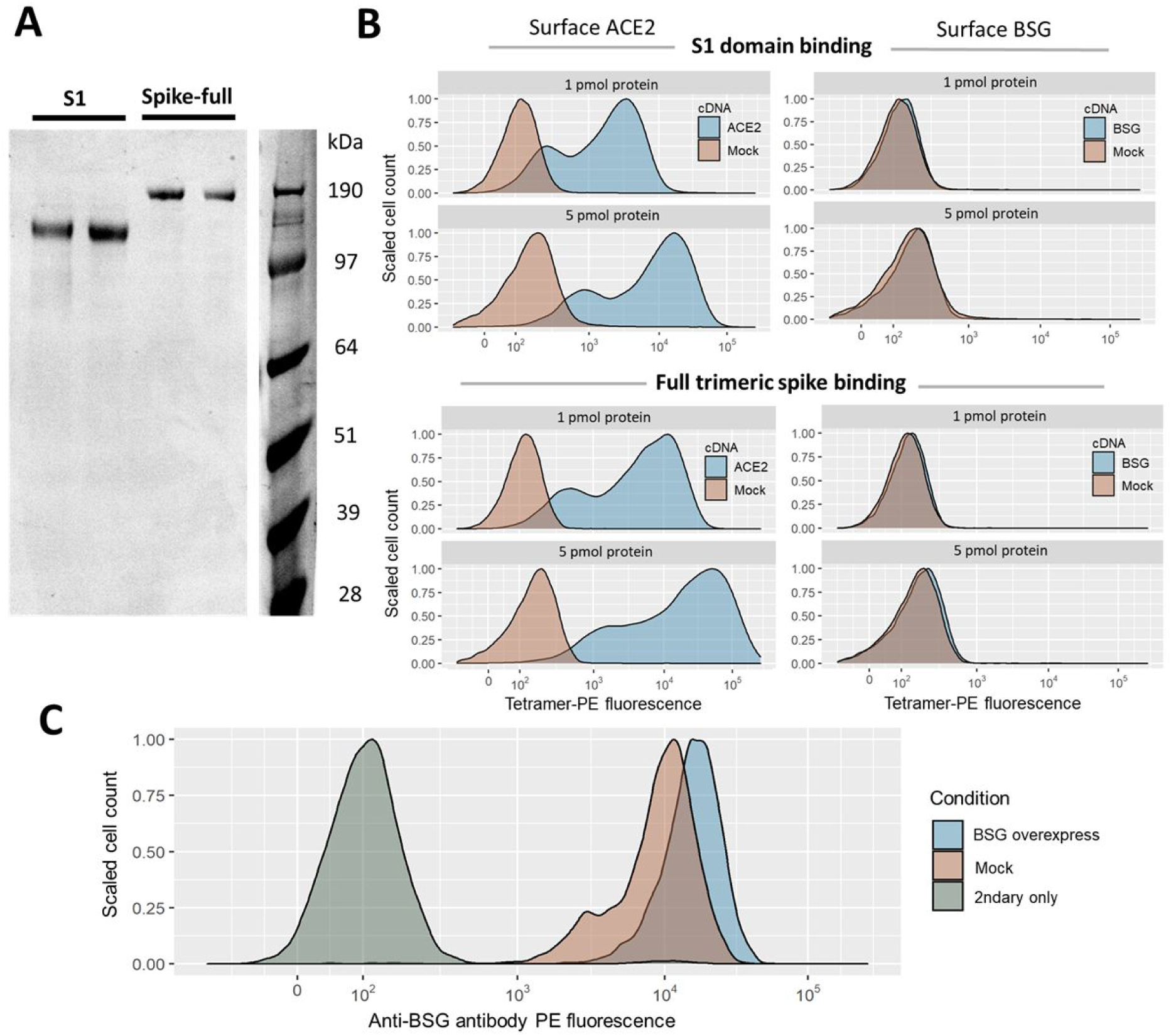
Gain of SARS-CoV-2 spike binding activity on human cells over-expressing ACE2 but not BSG. **A**. Expression and purification of the S1 domain and full ectodomain of the SARS-CoV-2 spike protein produced in human cell lines. Two independent preparations of purified spike were resolved by SDS-PAGE under reducing conditions and stained with Coomassie blue dye. **B**. Cells transfected with cDNAs encoding ACE2 but not BSG bind highly avid fluorescent SARS-CoV-2 spike tetramers. Flow cytometry fluorescence distributions of cells stained with tetramers made of biotinylated spike protein either using the S1 domain (top panels) or the entire ectodomain (lower panels) clustered around phycoerythrin-conjugated streptavidin. The stained HEK293 cells were transfected with cDNA to overexpress either ACE2 (left) or BSG (right). Mock-transfected cells are shown in red. Similar behavior to the data shown was observed in three separate tests. **C**. Transfection with BSG cDNA leads to upregulation of cell-surface BSG. Surface basigin levels on HEK293 cells labeled with anti-human BSG monoclonal antibody. BSG levels are compared to a negative control of secondary-antibody only.

Next, we sought to leverage the high sensitivity of direct biochemical binding assays to determine if these methods could detect any traces of basigin binding. We have previously expressed the ectodomain of the basigin receptor in a functionally active form and used it to discover pathogen ligands including *Plasmodium falciparum* RH5^11,22–23^. In a HEK293 human cell line, we expressed recombinant forms of the extracellular domains of both the canonical isoforms of basigin (BSG) that contains two Ig-like domains and the alternate isoform which contains an additional Ig-like domain (BSG-long) (Figure 2A). To confirm our recombinant constructs were folded and biochemically active, we probed the basigin constructs with three different monoclonal antibodies known to bind native basigin at the cell surface^12,25^ in enzyme-linked immunosorbent assays (ELISAs). All antibodies specifically bound to both of our recombinant basigin isoforms but not a negative control construct of recombinant rat Cd4 tag (Figure 2B). The protein epitopes recognized by these antibodies retained their conformation in recombinant basigin but less so in basigin that is denatured by heat and reducing agent treatment, with each antibody having between 2-fold to >10-fold reduced immunoreactivity after treatment (Figure 2C). To determine significance we fit log-logistic dose-response models to each protein and antibody combination that had at least 2 replicates^26^, with Bonferroni-corrected p-values ranging from 0.02 (BSG and MEM-M6/1) to < 0.0001 (all others) when comparing denatured and control curves by F-tests.

**Figure 2.**
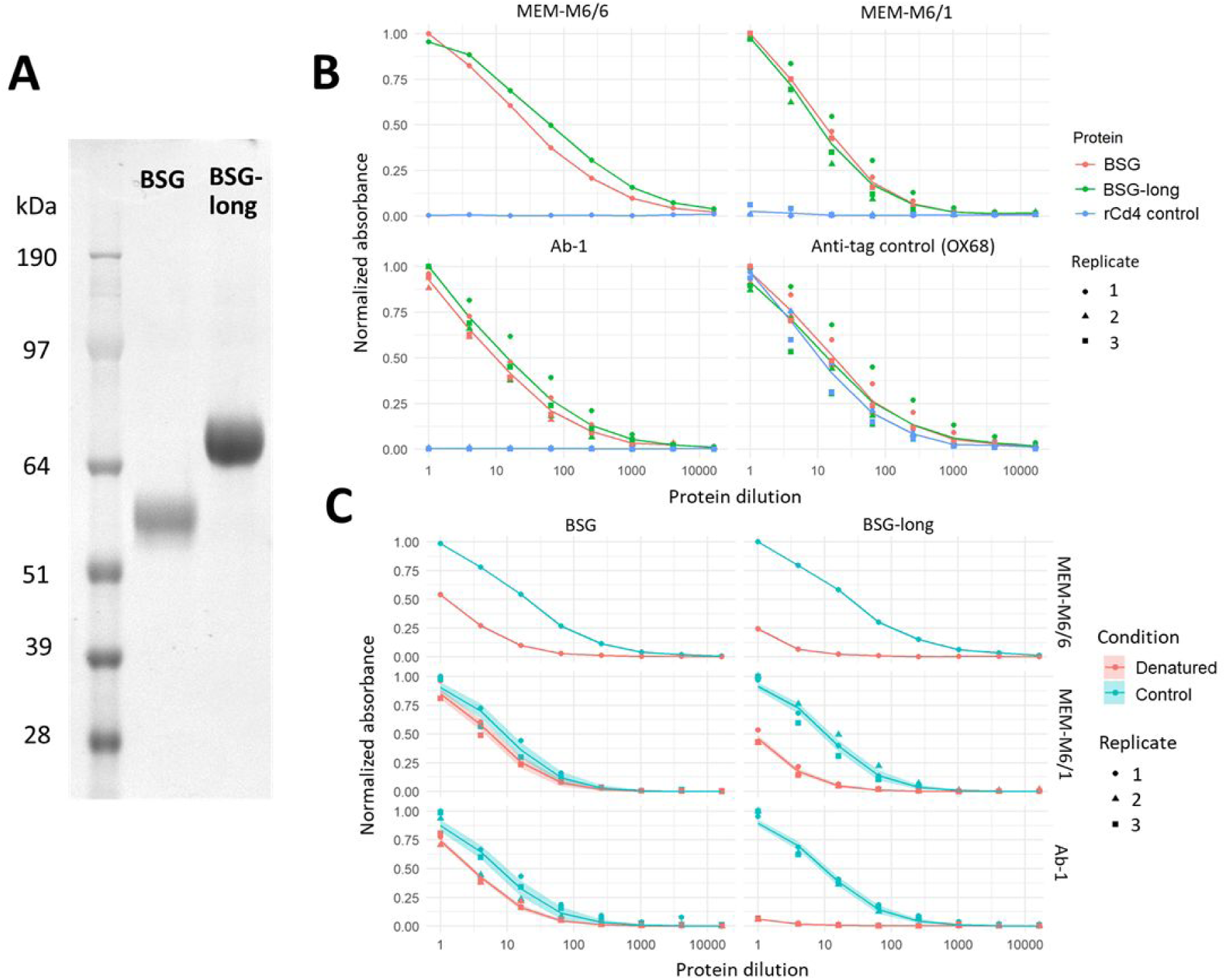
Recombinantly expressed basigin ectodomains retain biochemical activity. **A**. Expression and purification of two and three Ig-like domain forms of basigin. Proteins were resolved under reducing conditions by SDS-PAGE and stained with a Coomassie dye **B**. Recombinant basigin but not control proteins are recognized by anti-basigin monoclonal antibodies. ELISA dilution series of BSG and BSG-long recognized by 3 different monoclonal antibodies, and a control OX68 antibody against their tags. A negative control of a recombinant Cd4 tag is included for each antibody. **C**. Recombinant basigin retains folded conformation of epitopes recognized by three different monoclonal antibodies. ELISA dilution curves comparing unmodified basigin to protein treated with heat and reducing agent. Three replicates were performed for all ELISA curves except MEM-M6/6, for which only a single trial was done. Dose response curve model fit lines are superimposed on the data points, with shading indicating the 95% confidence bounds of the models.

With the functionality of our constructs quality-tested, we performed a plate-based binding assay^27^ that uses the avidity gains of multimerized proteins to detect even highly transient protein-protein interactions^28^. The SARS-CoV-2 spike proteins and ACE2 gave clear binding signals in both binding orientations as plate-bound baits and reporter-linked preys, yet no signals were observed for either BSG isoform against either spike construct (Figure 3A). By contrast the known interaction between BSG and *Plasmodium falciparum* RH5 was readily detected, as was a control low-affinity interaction between human CD200 and CD200R. Notably these interactions have similar or even weaker affinity than reported for the BSG-spike interaction^11,29^. Finally, in response to recent reports of a mutation in the SARS-CoV-2 spike that is rapidly displacing the reference sequence^30,31^, we also checked whether the D614G variant of the spike could bind BSG; again, we could not detect any interaction (Figure 3B). In all configurations the signal from BSG binding spike protein was indistinguishable from the background of non-interacting protein pairs and significantly below the known interaction pairs (Welch’s t-test of ACE2-spike interactions vs BSG-spike interactions p = 0.0002) (Figure 3C).

**Figure 3.**
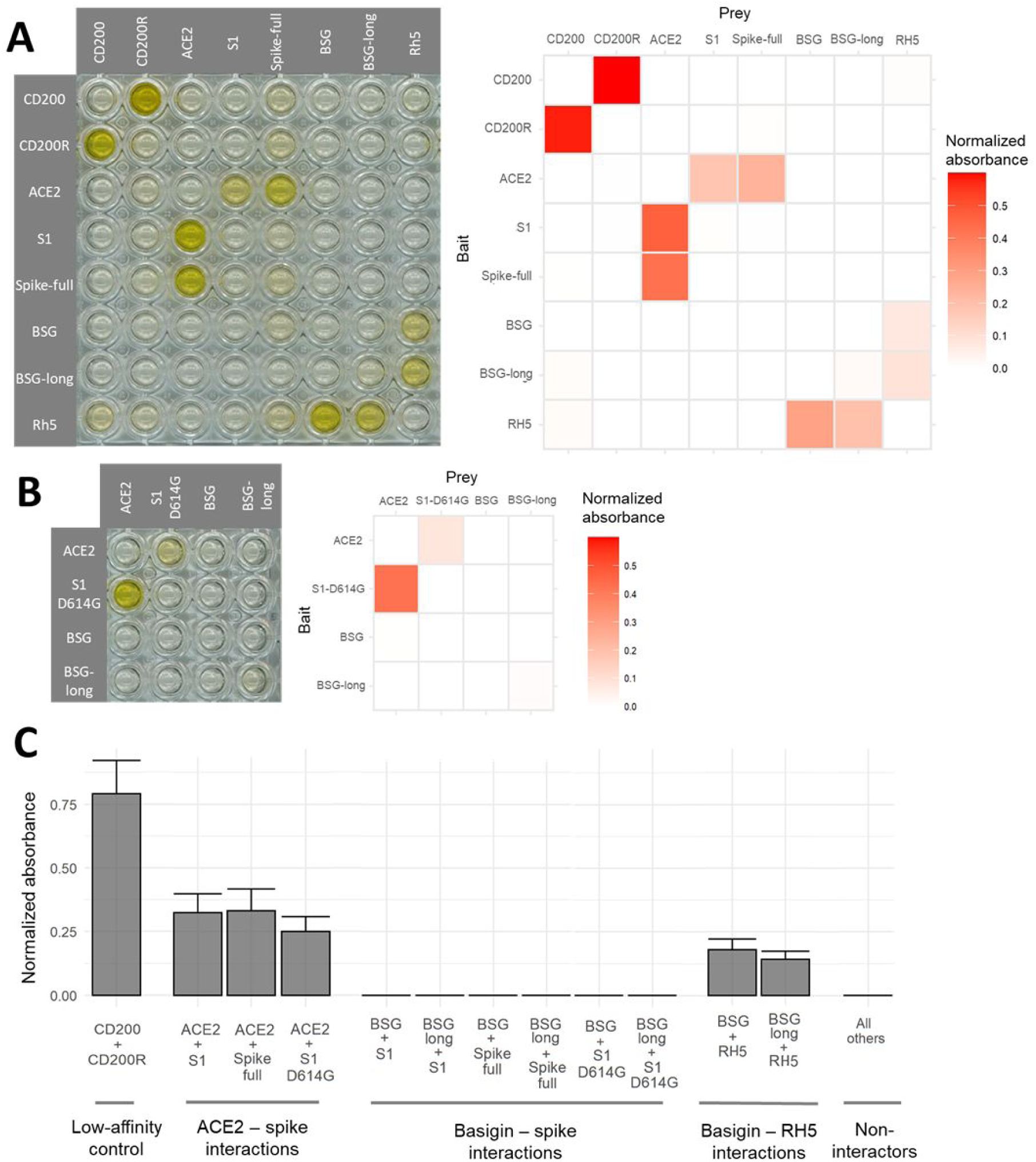
Sensitive assays designed to detect extracellular protein interactions do not detect a direct interaction between human basigin and the SARS-CoV-2 spike protein. **A**. No signs of spike-basigin binding in an avidity-based protein interaction assay testing a matrix of recombinant baits immobilized to streptavidin-coated plates (rows) against preys clustered around HRP-conjugated streptavidin (columns). An example raw screening plate (left) is shown alongside background-corrected absorbance values averaged across two replicates. **B**. The emerging D614G mutant variant of the SARS-CoV-2 spike also does not bind basigin. Binding matrix including the common D614G variant of the SARS-CoV-2 spike protein instead of the reference sequence. **C**. Spike binding to basigin is consistently undetectable compared to other control interactions. Binding signal averaged across bait and prey orientations for known interacting protein pairs, the basigin-spike pairs, and all other pairs. Error bars represent standard deviation from the mean (n = 2, except “All others” n = 58).

## Discussion

Identifying the host receptors which a virus can recognize is an important step in mechanistically explaining viral infection, and can offer insight in a virus’ cellular tropism and factors influencing susceptibility. Despite the importance of determining precisely which entry receptors SARS-CoV-2 uses to infect human cells, there remains considerable uncertainty amid multiple claimed viral receptors with variable qualities of data behind them^9,32–35^. We investigated one of the most prominent claims among these, that human BSG acts as an alternate receptor for the virus to interact with, which has been the topic of several studies, news and review articles, and a clinical trial^8,14,15,36–39^. Our access to established tools and reagents from previous work studying BSG’s role as a host receptor in *Plasmodium* infection allowed us to rapidly investigate BSG as a SARS-CoV-2 receptor. Despite validating the functionality of all our reagents, we were unable to detect any binding in biochemical or cell-based assays for either common BSG isoform or either configuration or allele of the SARS-CoV-2 spike protein.

BSG is highly expressed on many cell types throughout the body, including activated lymphocytes and red blood cells, forming the basis of the Ok blood grouping system^41^. Notably, SARS-CoV-2 has not been found to enter red blood cells^42^. However, the possibility that BSG could act as an accessory binding receptor for the virus has been speculated in several publications to possibly explain in part the link between SARS-CoV-2 infection and hematological symptoms in patients^38,39,43,44^. Our data suggest this hypothesis should be treated cautiously. Similarly, if our negative findings are replicated, it would necessitate a re-interpretation of the clinical trial involving injections of anti-BSG monoclonal antibodies, as any patient benefit would be more likely explained by alternative hypotheses such as immune modulation as opposed to direct blockage of viral invasion through BSG. Hypotheses relying on BSG binding to explain viral tropism may also need closer reconsideration^14^.

Although our findings were negative, they nevertheless carry important potential implications to both our understanding of the basic biology of SARS-CoV-2 and efforts to translate knowledge of the virus’ host receptors into therapeutics. We encourage greater study in confirming the mechanisms that have been proposed, not just for BSG but also for the multiple other putative viral receptors, so as to resolve the uncertainty around whether SARS-CoV-2 utilizes any receptors beyond ACE2 during infection.

## Materials and Methods

### Expression construct design

cDNA expression constructs were taken from a previously-assembled library of full-length human cDNAs in human expression vectors^24^. The BSG construct was cloned from a copy (Origene #RG203894) of the canonical 2-domain isoform of BSG (NM_198589.1), while the ACE2 cDNA (NM_021804.2) was expressed from a similar expression vector utilizing a CMV promoter (Geneocopia #EX-U1285-M02). The recombinant human BSG ectodomain constructs have been previously described^11,23^ and span from M1-L206 (BSG) or M1-L322 (BSG-long). The extracellular domain truncations for the SARS-CoV-2 spike proteins have also been previously published^17^. The S1 domain was defined as spanning Q14-Y647, while the full spike ectomain spanned Q14-K1211. The endogenous viral signal peptide was replaced with an efficient mouse antibody signal peptide^45^. As previously described, the full spike ectodomain was mutated at its polybasic protease cleavage site (682-685 RRAR to SGAG), had a proline stabilizing mutation introduced (986-987 KV to PP), and to mimic the natural trimerized structure of the spike had a foldon trimerization domain introduced at its C-terminus. The ACE2 ectodomain spanned M1-S740, retaining its endogenous signal peptide.

### Recombinant protein expression and purification

Human embryonic kidney (HEK)-293E cells were transiently transfected with polyethylenimine as previously described^46,47^. Per 100 mL of cells, 50 µg of plasmid was transfected along with 1 µg of a plasmid encoding the biotin-ligase BirA to direct biotinylation of the recombinant proteins^48^. Cells were grown in Freestyle Media (Life Technologies #12338018) supplemented with 100 µM D-biotin (Sigma #2031). Human proteins were incubated for 120 hours at 37°C, while spike proteins were shifted to 34°C and supplemented with 0.5% (m/v) tryptone N1 (OrganoTechnie #19553) 24 hours post-transfection and incubated a further 96 hours based on a published spike-specific optimized protocol^49^. After incubation, cell culture supernatants were harvested and passed through 0.22 µm filters. Purification was done using nickel-nitrilotriacetic acid (Ni-NTA) resins (Thermo Scientific #88221) that were pre-washed for 10 minutes in 2 washes of 25 mM imidazole (Sigma #I2399) phosphate buffer. Supernatants were mixed with pre-washed Ni-NTA resin overnight at 4°C, then washed three times with 25 mM imidazole phosphate buffer before eluting in 200 mM imidazole buffer. Purified proteins were analyzed on 4-12% gradient Bis-Tris gels (Invitrogen #NP0329) following denaturation for 10 minutes at 80°C in NuPAGE sample buffer (Invitrogen #NP0007, #NP0004). Across experimental replicates, independent batches of protein were used, with the exception of BSG and BSG-long for which a single batch was quality-tested and used in all subsequent experiments.

### Flow cytometry and tetramer binding assays

To generate transfected cells overexpressing cell-surface receptors, human embryonic kidney (HEK)-293E cells were seeded one day prior to transfection at a density of 2.5×10^5^ cells per mL in Freestyle Media (Life Technologies #12338018) supplemented with 10% heat-inactivated fetal bovine serum (FBS). Cells were transiently transfected with polyethylenimine as previously described^39,4^, except with double the ratio of DNA to cells. Then 48 hours after transfection, cells had culture media aspirated and were resuspended in 1 μM DAPI (Biolegend #422801) and incubated on ice for 5 minutes. Cells were then stained in u-bottom 96-well plates (Greiner #650161) with recombinant protein tetramers conjugated to phycoerythrin (PE). Tetramers were prepared by mixing 1 or 5 pmol of biotinylated protein monomer with 0.25 to 1.25 pmol of streptavidin-PE (Biolegend #405245) respectively and incubating for 2 hours at room temperature. Cells were spun down and resuspended in 100 μL of tetramer in a solution of 1% bovine serum albumin (BSA) and phosphate buffered-saline (PBS) supplemented with calcium and magnesium ions (Gibco #14040133). Cells and tetramers were incubated on ice for 45 minutes, washed in cold PBS, and stained for viability with DAPI (Invitrogen #D1306), and finally resuspended in the same 1% BSA PBS solution. Antibody staining was performed with a similar procedure, except during the first 30 minutes on ice cells were incubated with 30 μg/mL of monoclonal Ab-1 anti-BSG antibody^12^, then resuspended in 1:500 anti-human IgG antibody conjugated to Cy3 (Sigma #C2571). Fluorescence staining was measured by a BD Fortessa flow cytometer.

### Monoclonal antibody ELISAs

Streptavidin-coated 96-well plates (Nunc #436014) were pre-washed in 175 μL hepes-buffered saline (HBS) with 0.1% tween-20 (HBS-T), then blocked in 2% (m/v) bovine serum albumin (BSA, Sigma #A9647) in HBS for 1 hour at room temperature. In a separate 96-well plate, a 1:4 dilution series of biotinylated BSG or control protein was prepared in 2% BSA HBS, then 100 μL of the protein dilution transferred to the blocked streptavidin-coated 96-well plate. In the experiments to determine the protein’s sensitivity to heat and reduction treatment, one half of the protein sample was denatured by heating at 80°C for 10 minutes in the presence of 5% beta-mercaptoethanol. After capturing protein for 1 hour at room temperature, plates were washed three times with 150 μL HSB-T. Anti-human basigin monoclonal antibodies were added at the following concentrations: 1.7 μg/mL Ab-1 (Zenonos et al., formerly known as ch6D9)^12^, 2.2 μg/mL MEM-M6/1 (Abcam #ab666), and 1.3 μg/mL MEM-M6/6 (Abcam #ab119114)^25^. A control mouse anti-rat Cd4 domain 3+4 monoclonal antibody (OX68) against the tags of our recombinant proteins was used at a 1.6 μg/mL concentration. After 1 hour of incubation with the primary antibody and three HBS-T washes, secondary antibody was added as 1:7000 donkey anti-human IgG (Abcam #ab102407) for ch6D9 and for all other antibodies as 1:3500 goat anti-mouse IgG (Sigma #A9316). Both secondary antibodies were conjugated to alkaline phosphatase. After 45 minutes of incubation with the secondary antibody, the plates were washed again three times with HBS-T. A substrate of 60 μL 2 mg/mL para-Nitrophenylphosphate (Sigma #P4744) in diethanolamine buffer was added to each well to develop signal over 30 minutes. Absorbance was measured on a Tecan plate reader at 405 nm.

### Avidity-based binding assays

Biotinylated recombinant proteins were tetramerized around streptavidin-HRP (Pierce #21130) for 1 hour at room temperature to form reporter-linked preys. Per well of the assay plate, 0.1 pmol of recombinant monomer were added to 0.025 pmol of streptavidin to create a highly avid binding reagent. Streptavidin-coated 96-well plates (Nunc #436014) were pre-washed in 175 μL hepes-buffered saline (HBS) with 0.1% tween-20 (HBS-T), then blocked in 2% (m/v) bovine serum albumin (BSA, Sigma #A9647) in HBS for 1 hour at room temperature. Biotinylated baits were captured by adding 0.1 pmol of purified protein diluted in 2% BSA to each well of the plate. After incubating the baits for 2 hours at 4°C, plates were washed three times with 150 μL HBS-T. The pre-formed tetrameric preys were then added and the plate incubated for 1 hour at room temperature. The plate was finally washed twice in 150 μL HBS-T and once in 150 μL HBS before adding 60 μL 3,3′,5,5′-Tetramethylbenzidine (TMB) substrate (Sigma #T0440). After developing signal for 15 minutes at room temperature, the reaction was halted by the addition of 0.25 M HCl. Absorbance was measured on a Tecan plate reader at 405 nm.

### Data processing

For flow cytometry experiments, measurement events for analysis were gated on live singlet cells (based on DAPI and forward and side scatter profiles) using FlowJo version 9. No compensation was done because only a single fluorochrome was used for cell staining. Cytometry data was visualized using the CytoML package^50^ in R version 3.6.1. For plate-based experiments, raw absorbance values had background subtracted. Background for ELISAs was defined as the minimum absorbance of any well on the measured plate, and background for binding assays was defined as the median absorbance of each respective tetrameric prey. For better comparing replicates, these corrected absorbances were rescaled by min-max normalization so that the maximum absorbance on that replicate’s entire plate is defined as 1. Statistics on binding assay data were calculated using the t.test function in the R base stats package (version 3.6.1). Statistics on ELISA data were calculated by performing an F-test comparing a two-parameter log-logistic dose response model fitted to the ELISA data to a null model where both the denatured and control protein conditions were assumed to be identical. Model fitting and statistical procedures were done using the drc package in R as previously described^26^.

## Acknowledgements

We thank Thomas Crozier and Paul Lehner for helpful discussions.

## Author contributions

JS performed all experiments, analyzed all data, and wrote the manuscript. GJW supervised the research and edited the manuscript.

